# Effect of joint velocity and pre-activation on the torque-fascicle length relationship of the vastus lateralis

**DOI:** 10.64898/2026.06.23.734014

**Authors:** Tristan Tallio, Antoine Nordez, Lucie Lecarpentier, Sylvain Dorel

**Affiliations:** Nantes Université, Motricité - Interactions - Performance, MIP, UR 4334, F-44000, Nantes, France; Institut Universitaire de France (IUF), Paris, France

**Keywords:** Isokinetic contractions, Optimal angle, Optimal fascicle length, Pre-activation, Ultrasound

## Abstract

Fascicle operating length during dynamic tasks is often compared to the isometric torque-length relationship, but there is a lack of evidence regarding the influence of joint velocity on optimal fascicle length. Moreover, there is no consensus in the literature regarding the influence of contraction initiation (pre-activation or passive start), although it could alter the interaction between fascicles and the tendon. This study aimed to investigate the effect of joint velocity and pre-activation on the torque-angle and torque-length relationships of the vastus lateralis during mono-articular isokinetic knee extensions. Twenty-one participants performed isometric, isokinetic (50°.s^-1^ to 450°.s^-1^), and isokinetic knee extensions with maximal isometric or eccentric pre-activation at 100°.s^-1^ and 300°.s^-1^. Torque, joint angle, fascicle length, and electromyographic activity of the quadriceps femoris muscles were recorded during contractions and then used to model the torque-angle and torque-length relationships. We were able to successfully fit the torque-angle and torque-length relationships (R²=0.93 and R²=0.92, respectively). A main effect of velocity was detected regarding the optimal angle (p<0.05), but no significant change was observed for the optimal fascicle length. Isometric pre-activation induced a reduction in maximal torque production compared with eccentric pre-activation and passive conditions at both isokinetic velocities (p<0.001), with no change in muscle activity. Our results suggest that muscle-tendon interactions may permit a dissimilar behavior between the torque-angle and the torque-fascicle length relationships. The reduction in torque following isometric pre-activation may be related to a contraction history-dependent phenomenon.

**NEW & NOTEWORTHY:** We demonstrated that, at a given joint angle, increasing velocity altered fascicle operating length without shifting optimal fascicle length, likely because of muscle-tendon interactions. We also showed that maximal isometric pre-activation before a concentric contraction reduced mean and maximal torque during the isokinetic phase compared with eccentric pre-activation or no pre-activation. This effect may be linked to contraction history, since muscle activity did not differ between conditions.

## INTRODUCTION

The isometric torque-angle relationship is frequently used in single-joint movements (Noorkoiv et al., 2015; Wisdish et al., 2025), as a proxy for the muscle force-length relationship (Kellis & Blazevich, 2022). Indeed, the joint angle at which the maximal torque is produced (i.e. the optimal angle - Oranchuk et al., 2020) is considered to reflect the optimal muscle length for maximal force-generating capacity of the muscles involved (Fukashiro et al., 2006).

In dynamic conditions, beyond the direct effect of contraction velocity on force production, joint velocity changes the shape of this torque-angle relationship (Hahn et al., 2014). Several studies have investigated the three-dimensional relationship between torque, angle and joint velocity in the knee extensors (Anderson et al., 2007; Kawakami et al., 2002; Thorstensson et al., 1976). By assessing isokinetic knee extension torque at different velocities, they highlighted a shift in the optimal angle with increasing velocity (i.e. occurring in a more extended knee position). Kawakami et al. (2002) found a match between this shift and the elongation of the series elastic component, and speculated that the change in optimal angle might be attributed to the compliance of the series elastic component. Hence, they suggested that the optimal length is decoupled from the optimal angle, and remains unchanged with increasing joint velocity. However, to our knowledge, no previous study has confirmed this. Additionally, torque-angle and torque-length relationships are classically used as reference profiles to interpret the operating joint angles and muscle lengths during functional tasks, with the aim of estimating force-production capacity throughout the movement (Bohm et al., 2018; Nikolaidou et al., 2017). However, given the possible effect of velocity, it would be preferable to assess these relationships during dynamic contractions. Few studies have investigated the torque-angle relationship during dynamic contractions; however, these investigations were either conducted on different muscle groups at very slow angular velocities (Brockett et al., 2001; Guex et al., 2016) or performed during multi-joint movements (Hahn et al., 2014). More generally, no previous study has investigated the three-dimensional relationship between joint angle, joint velocity and torque through the prism of muscle fascicle length.

From a methodological point of view, the general assumption is that muscle contraction is maximal over the whole range of motion during isokinetic contractions, but there is no consensus in the literature about how the contraction should be initiated. In some studies, participants performed isokinetic contractions starting from a passive state and contracted “as hard as possible” to initiate the movement (Drazan et al., 2019; Hahn et al., 2014; Noorkoiv et al., 2015; Oranchuk et al., 2020). Others suggested the use of isometric or eccentric pre-activation prior to the measured concentric isokinetic contraction (Brown et al., 2016; Forrester et al., 2011; Germano Maciel et al., 2020; Pain et al., 2013). Pre-activation, also referred to as the “active state” (Bobbert & Casius, 2005), is a muscle contraction prior to movement considered as a way to i) improve muscle force during dynamic contractions, by minimizing incomplete muscle activation (Brito Fontana et al., 2014; Finni et al., 2003), and ii) reduce muscle-tendon unit slackness (Ichinose et al., 2000). Thus, a change in the optimal angle to a more flexed knee-joint angle may occur in contractions with pre-activity. Moreover, pre-activation might change the dynamic interactions between fascicles and tendons, inducing a reduction in fascicle length (Beaumatin et al., 2018; Goecking et al., 2024) and an increase in tendon length (Beaumatin et al., 2018). To our knowledge, the relationship between these changes in the muscle-tendon complex and the modifications in muscle force production has not been previously explored.

Therefore, this study aimed to analyze the effects of knee-joint velocity on maximal torque-angle and torque-length relationships at the level of the vastus lateralis fascicles. The second objective was to investigate how pre-activation enhances torque production during the subsequent dynamic contraction and modifies these relationships. To this end, vastus lateralis fascicle length was measured during maximal isokinetic knee extensions performed at multiple joint velocities (i.e. from 50°.s^-1^ to 450°.s^-1^) with an initially passive state, and at two additional joint velocities (i.e. 100°.s^-1^ and 300°.s^-1^) with eccentric or isometric pre-activation. We hypothesized that the change in optimal angle observed with increasing joint velocity would not be associated with a change in the optimal fascicle length, due to the dynamic interaction between fascicles and tendinous structures (Kawakami et al., 2002). We also expected an increase in torque production with eccentric and isometric pre-activation, accompanied by a modification of the optimal fascicle length caused by the initial shortening of fascicles (Beaumatin et al., 2018; Goecking et al., 2024).

## METHOD

### Participants

Twenty-one physically active sport science students (7 females, 14 males) volunteered to participate in this study (age 20.9 ± 2.3 years, height 177.0 ± 11.1 cm, body mass 68.8 ± 13.4 kg). They were free of any musculoskeletal injuries of the lower limbs for at least one year. Participants were informed about the nature, aims and risks of this study, and gave their written consent to participate. This study was approved by the local Institutional Review Board of the University (CEDIS #22042024-1).

### Experimental setup

#### Ultrasound

Two ultrasound systems (SuperSonic Imagine, v12, Aix-en-Provence, France) were used to visualize the vastus lateralis (VL) fascicles using ultrafast sequences (plane waves). The frequency was set at 100 Hz, 500 Hz or 1000 Hz, depending on the duration of the task. Dual probes were used to measure the entire length of the vastus lateralis fascicles in order to prevent the need for extrapolation (Brennan et al., 2017). The probes (15-4 MHz, 55 mm, Vermon, Tours, France) were placed in series with the help of a custom 3D-printed holder, and fixed on the right thigh at approximately 50% of the muscle length. The imaging plane was adjusted to observe fascicles on the two probes and to avoid blood vessels (Bolsterlee et al., 2016). The probes were then attached to the thigh using double-sided tape and straps.

#### Dynamometer

Participants were seated on an isokinetic dynamometer (Con-Trex MJ, CMV AG, Dübendorf, Switzerland) with the hip flexed at 80°. The right knee joint center (i.e. defined as the femur’s lateral epicondyle) was aligned with the dynamometer axis of rotation, and a pad was placed between the shank and the dynamometer arm. The left leg remained passive in a flexed position during measurements (i.e. ∼90°). Participants were fastened to the chair with seatbelts, and the thigh and shank were attached to the chair and to the dynamometer’s arm, respectively, to avoid any movement. Torque and angle were recorded at 5 kHz using an analog-to-digital (A/D) converter (LockLab, Vicon Motion System Ltf, Oxford, UK).

#### Motion capture

Lower-limb movements were recorded using a 14-camera motion-capturing system (Vicon Motion System, UK). Twenty-six reflective markers (diameter of 14 mm) were placed on the pelvis, right thigh, right shank and right foot to reconstruct body segments. Markers were assigned to anatomical landmarks (anterior and posterior superior iliac spines, iliac crests, femoral medial and lateral epicondyles, anterior tibial tuberosity, medial and lateral malleolus, 1^st^ and 5^th^ metatarsal bone and hallux), and to technical landmarks forming clusters (4 on the thigh, 4 on the shank, 2 on the foot).

#### Electromyographic measurements

Myoelectric activity of the VL, rectus femoris (RF) and vastus medialis (VM) muscles was recorded using surface electromyography (PicoEMG, Cometa, Bareggio, Italy). The sampling rate was set at 2 kHz and data was synchronized to the A/D converter. After skin preparation, the electrodes were placed on the belly of the three muscles according to the SENIAM guidelines (http://www.seniam.org).

#### Synchronisation

Torque and angle data from the dynamometer were recorded using the A/D converter. The EMG software was synchronized with the A/D converter so that recording began simultaneously with the motion capture measurements. Ultrasound data were synchronized with all devices using a pedal, which started ultrasound recording and sent a Dirac signal to the A/D converter. For data processing, all measurements were cropped from the onset of the Dirac signal and resampled to match the A/D frequency (i.e. 5 kHz).

### Experimental design

Following a standardized warm-up on the dynamometer, participants were asked to randomly perform maximal voluntary isometric contractions (MVIC) at 45°, 60°, 70°, 80°, 95° and 110° knee angles (0° corresponding to full knee extension) for approximately two seconds. In addition, they performed maximal isokinetic contractions at 50°.s^-1^, 100°.s^-1^, 200°.s^-1^, 300°.s^-1^ and 450°.s^-1^ in a randomized order. The starting knee joint angle was set at 120° and the final knee extension angle at 15°. Participants were asked to be fully relaxed before contraction to avoid any pre-activation. For all isometric and isokinetic contractions, participants were instructed to “push as hard as possible”. Participants were then asked to perform maximal isokinetic contractions at 100°.s^-1^ and 300°.s^-1^ under two pre-activation conditions (i.e. isometric and eccentric). For the isometric pre-activation condition, they performed an MVIC at 120° for approximately one second. For the eccentric pre-activation condition, they performed a maximal eccentric contraction over the same range of motion at 200°.s^-1^ and then continued to “push as hard as possible” throughout the subsequent concentric contraction phase. Two trials were performed for each condition, and a two-minute rest period was provided between trials. Finally, a passive trial at 5°.s^-1^ was performed across the entire range of motion. In all conditions, we measured active fascicle length, knee angle via motion-capture analysis, joint torque, and EMG activity.

### Data processing

#### Ultrasound

Images of the two probes were combined using a custom-written Matlab script (The MathWorks, Natick, MA, USA), creating a black space corresponding to the gap size between the two acoustic bands (i.e. 16 mm) in the combined image (Figure 1). Fascicle length was then measured using a semi-automated software (Ultratrack; Farris & Lichtwark, 2016). The superficial aponeurosis was defined primarily on the right image, and the deep aponeurosis on the left image. Fascicles were tracked only on the left image. The right image was used to measure the superficial aponeurosis and improve the linear extrapolation of fascicles (Werkhausen et al., 2022). For the isometric contractions, ultrasound data were filtered with a dual 10 Hz low-pass (2^nd^ order) Butterworth filter, and the mean fascicle length at peak torque was used for analysis. During isokinetic contractions, ultrasound data were filtered with a dual low-pass (2^nd^ order) Butterworth filter at 5 Hz, 10 Hz, 15 Hz, 20 Hz, and 25 Hz, for joint velocities set at 50°.s^-1^, 100°.s^-1^, 200°.s^-1^, 300°.s^-1^, and 450°.s^-1^, respectively (Hauraix et al., 2017).

**Figure 1:**
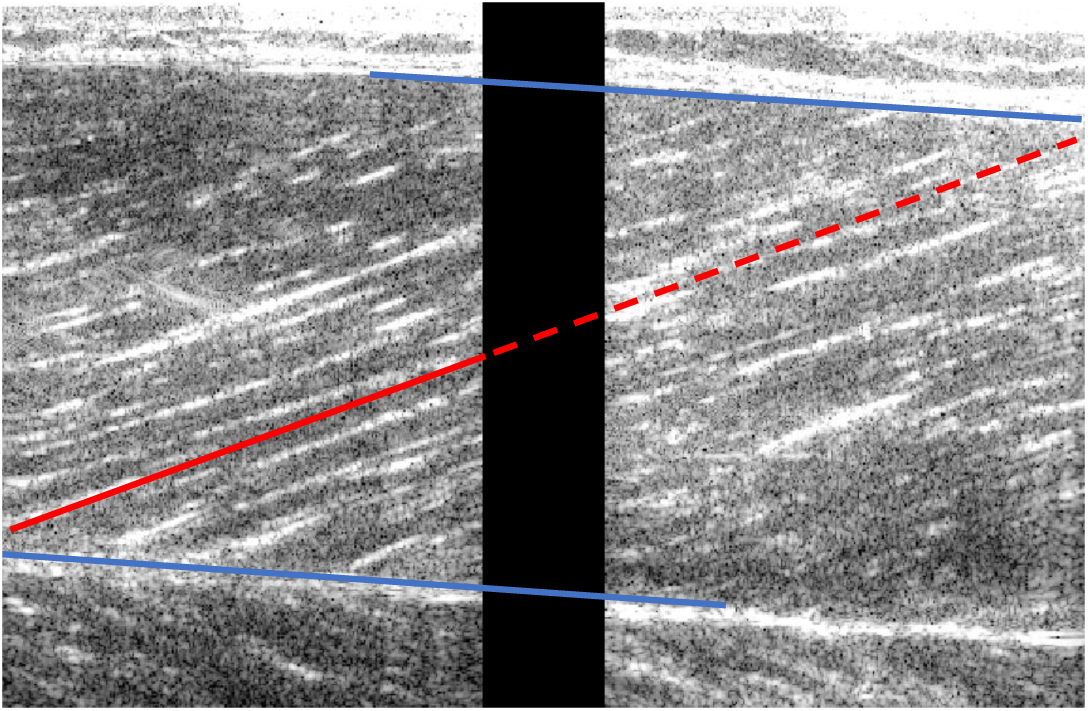
Example image of combined scans of vastus lateralis, showing the superficial and deep aponeurosis (blue), and the fascicle (red). The solid lines represent measured data, whereas the dashed line represents extrapolated data.

#### Mechanical data and joint kinematics

Knee joint angle under all conditions was calculated from motion-capture data. Three 3D local coordinate systems were constructed using three markers placed on the thigh, the femoral epicondyles, and the shank. Euler rotation matrices were applied to calculate the angle between the thigh and shank local coordinate systems, using the third axis as the axis of rotation. θ was defined as the knee joint angle.

During all contractions, torque was corrected for the gravitational forces of the leg, foot, and dynamometer arm for each trial to obtain the active torque (T_active_). During isometric contractions, torque and angle data were filtered with a dual 10 Hz low-pass (2^nd^ order) Butterworth filter. Peak T_active_ was calculated using a moving 200-ms averaging window. During isokinetic contractions, with and without pre-activation, torque and angle data were filtered with dual low-pass (2^nd^ order) Butterworth filter with cut-off frequencies of 5 Hz, 7.5 Hz, 10 Hz, 12.5 Hz, and 15 Hz, for joint velocities set at 50°.s^-1^, 100°.s^-1^, 200°.s^-1^, 300°.s^-1^, and 450°.s^-1^, respectively (Hauraix et al., 2017). Torque was subsequently corrected for inertial effects by accounting for joint acceleration (i.e. second derivative of the knee angle) and the masses of the leg, foot and dynamometer arm.

#### Torque-angle and torque-fascicle length modelling

Torque-angle and torque-fascicle length relationships were modelled using the equation proposed by Hoffman et al. (2012 - Eq. 1; Eq. 2). Knee extension active torque, angle, and fascicle length data were used to fit individual torque-angle and torque-length relationships using nonlinear least-square optimization (the *lsqcurvefit* function in Matlab):

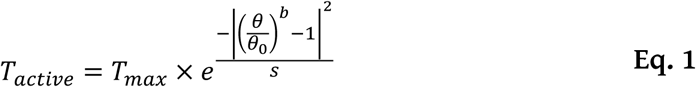

Where *T_active_* represents the active torque, *T_max_* the theoretical maximal active torque, *θ* the angle, *θ_0_* the optimal angle, *b* the skewness and *s* the width of the curve.

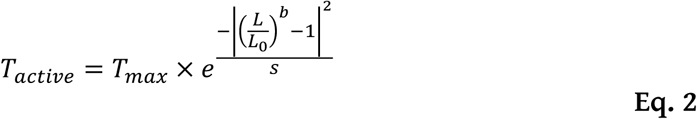

Where *T_active_* represents the active torque, *T_max_* the theoretical maximal active torque, *L* the fascicle length, *L_0_* the optimal fascicle length, *b* the skewness and *s* the width of the curve.

For the isometric conditions, the modeling analysis considered the peak T_active_ values of the five contractions performed at the different knee angles. For the isokinetic conditions, the modeling procedure included only data collected during the period of constant angular velocity (i.e. when acceleration was negligible), to avoid a confounding effect of joint velocity across the range of motion, as illustrated in Figure 2. Absolute values of θ_0,_ L_0_, and T_max_ for each velocity condition were determined from the fitted curves and were also expressed relative to the optimal joint angle and optimal length obtained under isometric conditions.

**Figure 2:**
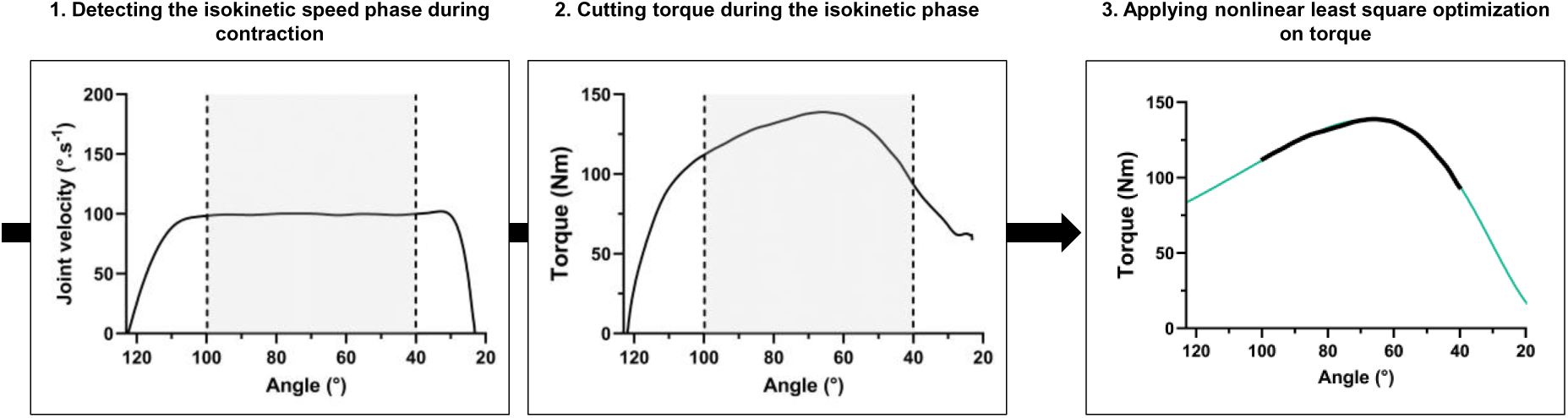
Example of the data-processing for modeling. The isokinetic portion of the contraction was identified (gray box between the dashed lines) and used to isolate the corresponding torque data (gray box between the dashed lines). The Hoffman equation (Eq. 1) was then applied to the retained data (bold black line) in a nonlinear least-square optimization function, resulting in the torque-angle relationship (green curve).

#### Electromyographic measurements

Raw EMG data were filtered using a 20–500 Hz bandpass (2^nd^ order) Butterworth filter, according to the recommendations of McManus et al. (2021), and then rectified. The data were subsequently smoothed using a dual low-pass (2^nd^ order) Butterworth filter with a cut-off frequency ranging from 10 to 25 Hz, depending on movement velocity. The maximal EMG activity level was detected using a moving time window adapted to joint velocity (i.e. from 100 ms for isometric contractions, to 35 ms for contractions performed at 450°.s^-1^). All EMG signals were then normalized to the maximal EMG level recorded across all contractions during the experimental session. The mean relative EMG activity level throughout the entire contraction was used to assess the effect of velocity, and the relative EMG activity level at the onset of movement was used for the pre-activation analysis.

### Statistical analysis

Normality was confirmed with a Shapiro-Wilk test. Data sphericity was assessed using Mauchly’s test, and corrected with the Greenhouse-Geisser coefficient when applicable. One-way repeated-measure analyses of variances (ANOVAs) were conducted separately to test the main effect of joint velocity and the effect of pre-activation. To examine the effect of joint velocity, ANOVAs were conducted on mean relative EMG activity, as well as on T_max_, θ_0_, and L_0_ derived from the torque-angle and torque-length models. To assess the effect of pre-activation, ANOVAs were conducted on the following variables derived from the experimental data: torque, fascicle length, and EMG activity at the beginning of the contraction (1^st^ frame), as well as mean torque and EMG activity during the isokinetic phase. ANOVAs were also performed on T_max_, θ_0_, and L_0_ whenever modelling was possible. For all analyses, post-hoc pairwise comparisons with Bonferroni correction were performed when appropriate. The significance level was set at α = 0.05, and data are presented as means ± standard deviation.

## RESULTS

### Effect of joint velocity

#### Experimental data

Mean experimental torque-angle, joint velocity-angle, torque-fascicle length, and fascicle velocity-angle relationships are presented in Figure 3. Joint angle at the start (123.2 ± 6.1°) and at the end (30.3 ± 12.5°) of the contraction did not differ between conditions, nor did fascicle length at the start (11.2 ± 0.1 cm) and at the end (7.3 ± 0.1 cm) of the contraction. Visual inspection of the curves confirmed that joint velocity, as well as fascicle velocity, increased concomitantly from the condition at 50°.s^-1^ to 450°.s^-1^. The isokinetic phase retained for analysis was from 110° to 40° for 50°.s^-1^, 100° to 40° for 100°.s^-1^, 90° to 45° for 200°.s^-1^, 80° to 45° for 300°.s^-1^, and 80° to 60° for 450°.s^-1^. Joint velocity over the range of motion reached the targeted plateau for the 50°.s^-1^ to 300°.s^-1^ isokinetic conditions, but not at 450°.s^-1^, and fascicle velocity also reached a plateau for the 50°.s^-1^ to 300°.s^-1^ isokinetic conditions, but not at 450°.s^-1^, due to the shorter isokinetic phase. Torque decreased with increasing isokinetic velocity, but the mean relative EMG activity did not differ among the VL, VM, and RF muscles (p-values from 0.070 to 0.143). The mean VL EMG amplitude ranged from 38.11 ± 4.09% to 43.75 ± 9.65%, the mean VM EMG amplitude ranged from 39.16 ± 8.78% to 45.53 ± 7.93%, and the mean RF EMG activity ranged from 40.24 ± 6.17% to 46.03 ± 8.80%.

**Figure 3:**
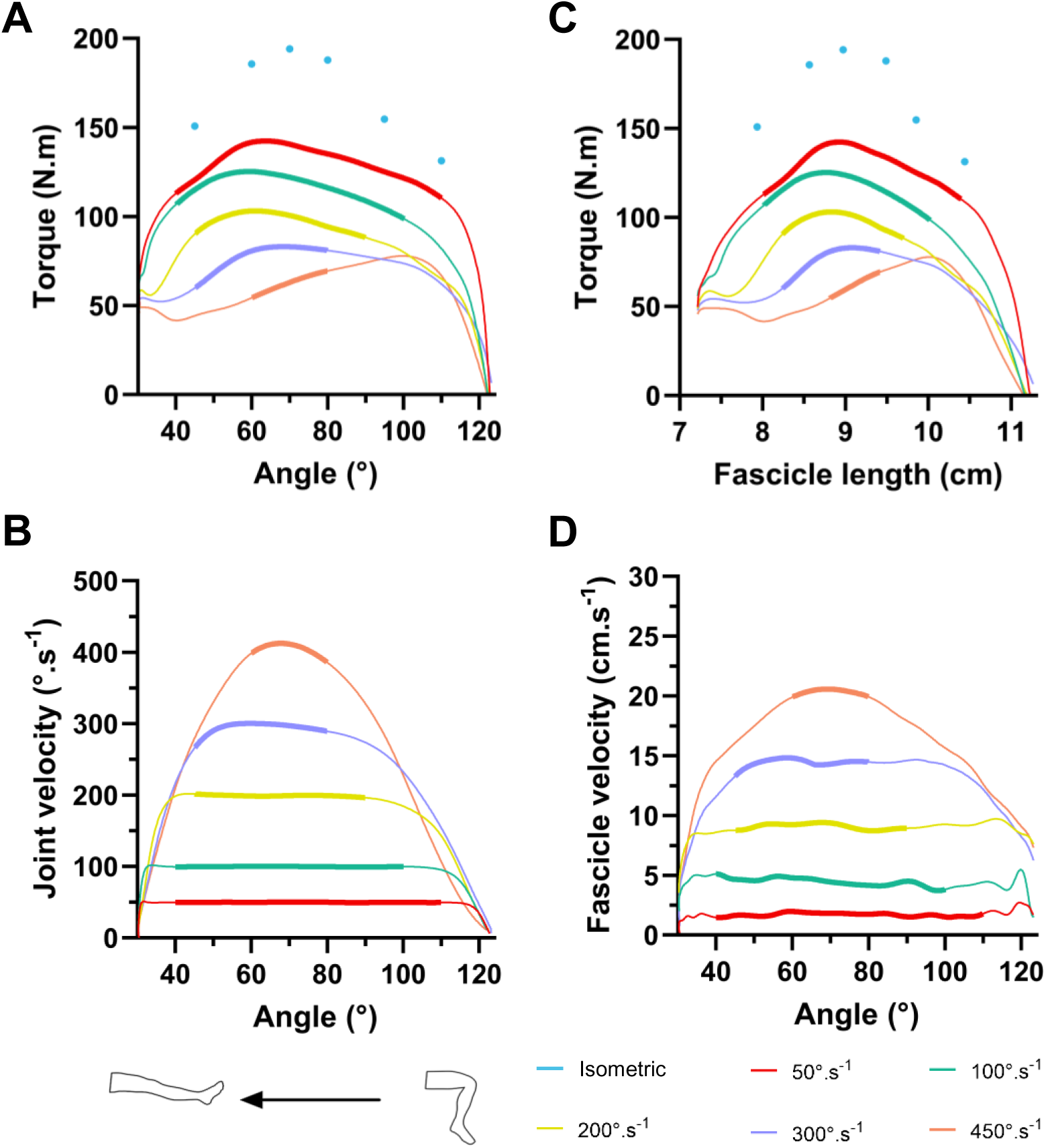
The evolution of torque as a function of joint angle (A) and fascicle length (C), and of joint velocity (B) and fascicle shortening velocity (D) as a function of joint angle during isometric and isokinetic contractions. Data were obtained during isometric contractions at 110°, 95°, 80°, 70°, 60°, and 45° of knee extension, and from 123° to 30° during isokinetic contractions. Bold lines represent data from the isokinetic portion used for the modeling analysis. Data presented are means; standard deviation are not shown for clarity. N=21.

#### Torque-angle and torque-length relationships modeling

Figure 3 shows the isokinetic phase retained for the modeling analysis (Figure 3B), and the associated torque values for the torque-angle (Figure 3A) and torque-length relationships (Figure 3C). In contrast to the isometric and slower velocity conditions, the pseudo-isokinetic phase in the 450°.s^-1^ condition was too short to display a curvilinear torque profile (Figure 3A and 3C). Hence, considering the lack of experimental data, torque-angle and torque-length relationships at this velocity could not be adequately and robustly modeled (see Supplementary Fig. S1), and will not be included further in the analysis. For the isometric, 50°.s^-1^, 100°.s^-1^, 200°.s^-1^, and 300°.s^-1^ conditions, torque-angle and torque-length relationships were well fitted by the model (Figure 4, Supplementary Fig. S1) with very high values of R²: a mean R² of 0.93 ± 0.09 for the torque-angle relationship (from 0.89 ± 0.12 at 50°.s^-1^ to 0.97 ± 0.03 at 300°.s^-1^, Figure 4), and a mean R² of 0.91 ± 0.10 for the torque-fascicle length relationship (from 0.87 ± 0.15 at 50°.s^-1^ to 0.95 ± 0.05 at 100°.s^-1^, Figure 4).

**Figure 4:**
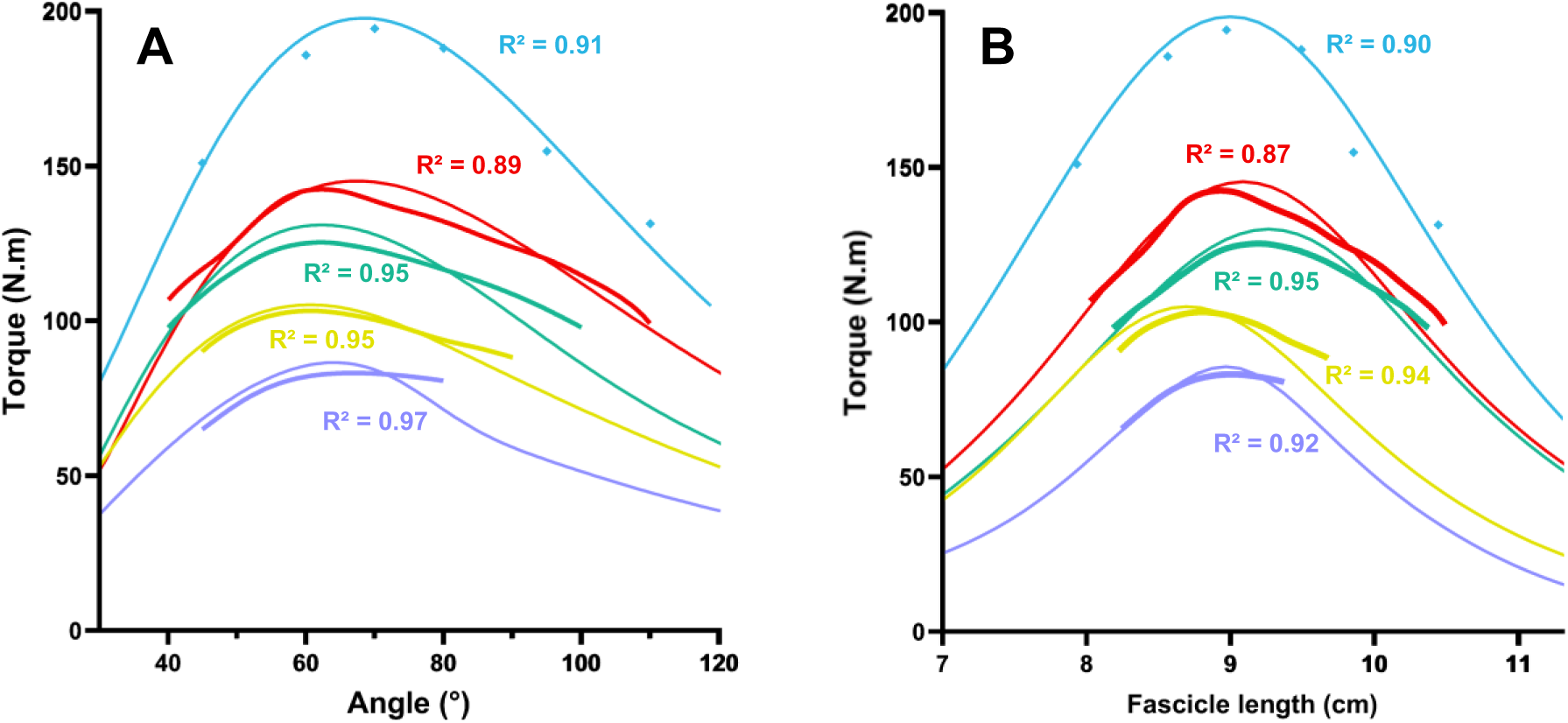
Mean torque-angle (A) and torque-length (B) relationships in isometric (light blue) and isokinetic conditions at 50°.s^-1^ (red), 100°.s^-1^ (green), 200°.s^-1^ (yellow) and 300°.s^-1^ (purple). Experimental data are represented by bold lines and squares, and fitted data by thin lines. Mean R² values for the modeling are presented for each condition.

Torque-angle and torque-length relationships normalized to θ_0_ or L_0_, and T_max_ are shown in Figure 5 (panels A and B, respectively). ANOVA revealed a main effect of joint velocity on T_max_ (p<0.001, Figure 5E). Maximal torque decreased with increasing speed, ranging from 199.0 ± 61.0 N.m in the isometric condition to 86.4 ± 26.2 N.m at 300°.s^-1^. Post-hoc analysis revealed significant differences between all conditions (p<0.001). ANOVA also revealed a main effect of joint velocity on absolute θ_0_ (p<0.05, Figure 5C), but no differences were found in post-hoc analysis (p-value ranging from 0.069 to 1.000). The absolute θ_0_ was reached between 68.5 ± 7.7° in the isometric condition and 60.6 ± 11.9° in the 200°.s^-1^ condition. No main effect of joint velocity was found for absolute L_0_ (p=0.071, Figure 5D). L_0_ ranged between 9.2 ± 1.5 cm in the 100°.s^-1^ condition and 8.8 ± 1.4 cm in the 200°.s^-1^ condition.

**Figure 5:**
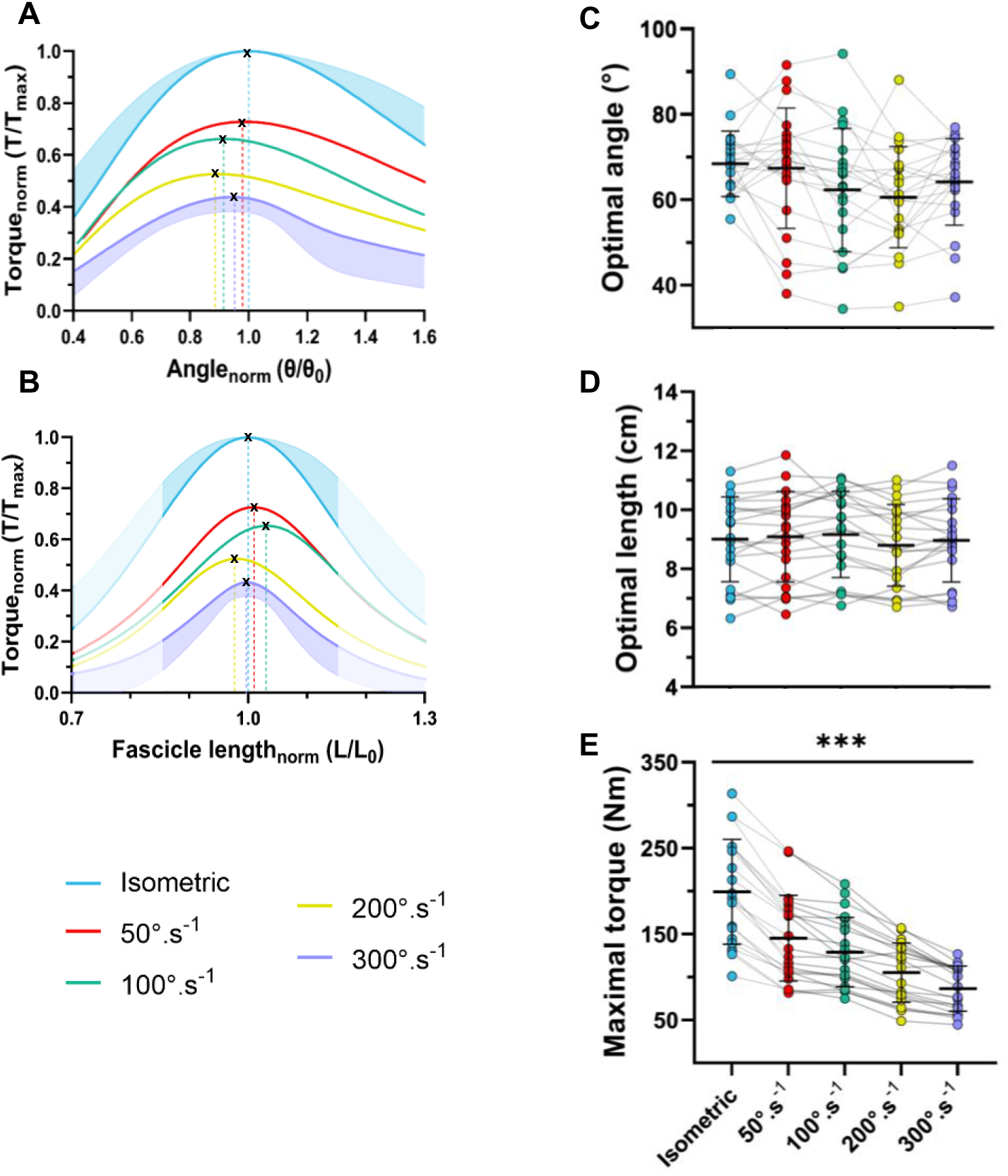
Modeling of the relative torque-angle (A) and relative torque-length (B) relationships during isometric, 50°.s^-1^, 100°.s^-1^, 200°.s^-1^, and 300°.s^-1^ conditions. Relative optimal torque, angle, and length were obtained by dividing the torque, angle, or fascicle length by the maximal torque, optimal angle, or optimal length of the isometric condition. Data are presented as means (solid lines), and only the standard deviations of the isometric and 300°.s^-1^ conditions are shown for clarity. Dashed lines represent projections of the optimal angle (A) and optimal length (B). For panels A and B, non-shaded data correspond to the range of data observed during experiments. Panels C, D, and E represent means and standard deviations (black) and individual data (colored dots) for the optimal angle, optimal fascicle length, and maximal torque, respectively. We found a main effect of velocity for the optimal angle and maximal torque, and *** indicates a significant difference between all conditions in post-hoc analysis (p<0.001). N=21.

### Effect of pre-activation

Results regarding the effects of pre-activation on torque, length, and joint angle are presented for 20 participants only, due to technical issues affecting one participant.

#### Experimental data

Figure 6 depicts torque as a function of joint angle and fascicle length, as well as joint and fascicle velocities as functions of joint angle, during the concentric contractions performed at 100°.s^-1^ (Figures 6A to 6D) and 300°.s^-1^ (Figures 6E to 6G) in each pre-activation condition. At 100°.s^-1^, ANOVA revealed a main effect of pre-activation on torque production at the beginning of the concentric contraction (p<0.001). Torque was significantly lower in NoPRE (2.6 ± 19.6 N.m) compared with PRE_iso_ (122.1 ± 37.7 N.m, p<0.001) and PRE_ecc_ (128.6 ± 48.3 N.m, p<0.001), but no difference was found between PRE_iso_ and PRE_ecc_ (p=0.425). ANOVA also revealed an effect of pre-activation on fascicle length at the start of the concentric contraction (p<0.001). Fascicle length was significantly longer in NoPRE (11.4 ± 1.7 cm) compared with PRE_iso_ (10.1 ± 1.6 cm, p<0.001) and PRE_ecc_ (10.6 ± 1.5 cm, p<0.01), and in PRE_ecc_ compared with PRE_iso_ (p<0.01). The mean torque produced during the isokinetic phase was lower in PRE_iso_ (101.6 ± 37.8 N.m) compared with NoPRE (111.1 ± 37.5 N.m, p<0.05) and PRE_ecc_ (118.6 ± 38.5 N.m, p<0.001), while no significant difference was found between NoPRE and PRE_ecc_ (p=0.173).

**Figure 6:**
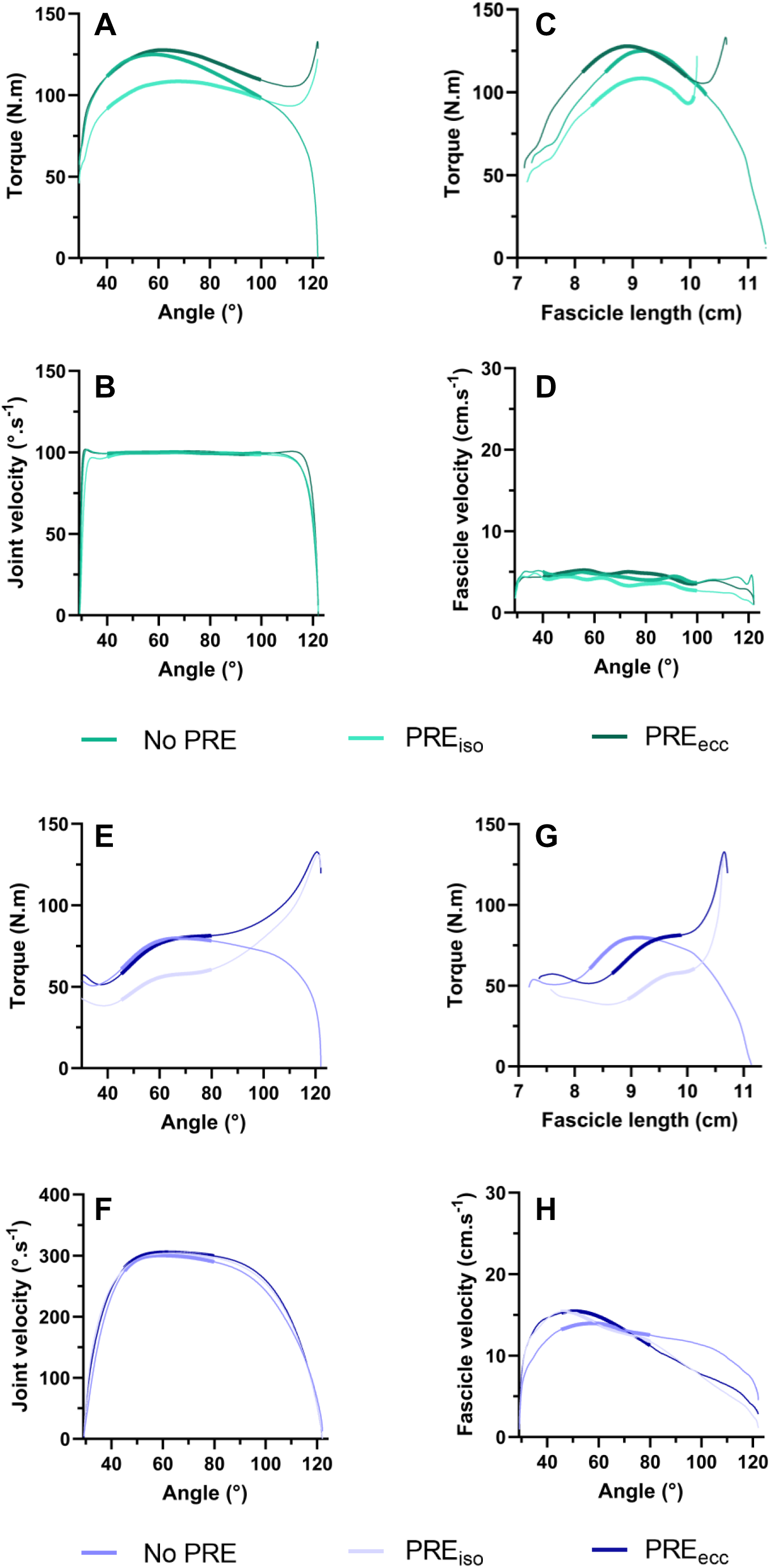
The evolution of torque as a function of joint angle (A and E) and fascicle length (C and G), and of joint velocity (B and F) and fascicle shortening velocity (D and H) as a function of joint angle during isokinetic contractions at 100°.s^-1^ (A to D) and 300°.s^-1^ (E to H). NoPRE corresponds to an isokinetic contraction alone, PRE_iso_ to an isokinetic contraction preceded by an isometric contraction, and PRE_ecc_ to an isokinetic contraction preceded by an eccentric contraction. Bold lines represent the experimental data used for the modeling analysis. Data are means; standard deviations are not shown for clarity. N=20.

At 300°.s^-1^, we found a main effect of pre-activation on torque production at the beginning of the concentric contraction (p<0.001). Torque was significantly lower in NoPRE (8.9 ± 25.2 N.m) compared with PRE_iso_ (123.1 ± 33.7 N.m, p<0.001) and PRE_ecc_ (119.6 ± 42.9 N.m, p<0.001), while no difference was found between PRE_iso_ and PRE_ecc_ (p=0.525). We also found a main effect of pre-activation on fascicle length at the start of the concentric contraction (p<0.01). Fascicle length was significantly longer in NoPRE (11.1 ± 1.7 cm) compared with PRE_iso_ (10.6 ± 1.7 cm, p<0.01) and PRE_ecc_ (10.7 ± 1.7 cm, p<0.05), and no difference was found between PRE_iso_ and PRE_ecc_ (p=0.578). The mean torque produced during the isokinetic phase was lower in PRE_iso_ (50.4 ± 20.6 N.m) compared with NoPRE (73.6 ± 27.5 N.m, p<0.001) and PRE_ecc_ (74.1 ± 29.0 N.m, p<0.001), while no significant difference was found between NoPRE and PRE_ecc_ (p=1.000).

Muscle activity of the VL, VM, and RF at the beginning of the concentric contraction and during the isokinetic phase at 100°.s^-1^ and 300°.s^-1^ is presented in Table 1. ANOVA revealed a main effect of pre-activation on the relative EMG amplitude at the beginning of the concentric contraction for the VL, VM, and RF (p<0.001 for all) at 100°.s^-1^. For the VL, the EMG amplitude was lower in NoPRE compared with PRE_iso_ (p<0.001) and PRE_ecc_ (p<0.001). Also, PRE_iso_ was higher than PRE_ecc_ (p<0.05). For the VM, the EMG amplitude was lower in NoPRE compared with PRE_iso_ (p<0.001) and PRE_ecc_ (p<0.05), and PRE_iso_ was higher than PRE_ecc_ (p<0.05). For the RF, the EMG amplitude was lower in NoPRE compared with PRE_iso_ (p<0.001) and PRE_ecc_ (p<0.001), while PRE_iso_ was not significantly different from PRE_ecc_ (p=0.134). Regarding the EMG activity during the isokinetic phase, no effect of pre-activation was found for the VL, VM, and RF (p=0.547, p=0.053, and p=0.797, respectively). For the 300°.s^-1^ condition, no effect of pre-activation was found on relative EMG activity at the beginning of the concentric contraction for the VL and VM (p=0.413 and p=0.215, respectively), but a significant effect was found for the RF (p<0.05), for which the EMG amplitude was lower in NoPRE compared with PRE_iso_ and PRE_ecc_ (p<0.05). No difference was found between PRE_iso_ and PRE_ecc_ (p=0.427). No effect of pre-activation was observed on mean EMG activity during the isokinetic phase for the VL, VM, and RF (p=0.319, p=0.626, and p=0.680, respectively).

**Table 1:** Mean muscle activation of the VL, VM, and RF at the start of and during the isokinetic phase of the contraction at 100°.s^-1^ and 300°.s^-1^. ^a^ and ^aa^ represent a main effect of pre-activation (p<0.05 and p<0.001, respectively), * and ** represent a significant difference compared with PRE_iso_ (p<0.05 and p<0.001, respectively), whereas # and ## represent a significant difference compared with PRE_ecc_ (p<0.05 and p<0.001, respectively). Data are presented as means ± standard deviations.

|  | Onset of concentric contraction |  |  | Isokinetic phase |  |  |
| --- | --- | --- | --- | --- | --- | --- |
| | NoPRE | $PRE_{iso}$ | $PRE_{ecc}$ | NoPRE | $PRE_{iso}$ | $PRE_{ecc}$ |
| **$100^{\circ}.s^{-1}$ condition** |  |  |  |  |  |  |
| **VL (%max) <sup>a</sup>** | 9.1 ± 12.9*## | 44.6 ± 15.6## | 33.6 ± 13.5 | 43.6 ± 7.8 | 42.9 ± 10.5 | 45.7 ± 10.2 |
| **VM (%max)<sup>a</sup>** | 14.3 ± 15.8*<sup>#</sup> | 48.4 ± 19.7<sup>##</sup> | 34.1 ± 12.0 | 45.3 ± 9.8 | 43.6 ± 12.0 | 49.8 ± 14.5 |
| **RF (%max)<sup>a</sup>** | 7.4 ± 11.0*<sup>##</sup> | 41.4 ± 16.8 | 32.4 ± 11.6 | 48.9 ± 8.6 | 49.2 ± 9.6 | 50.8 ± 11.5 |
| **300°.s<sup>-1</sup> condition** |  |  |  |  |  |  |
| **VL (%max)** | 26.4 ± 26.7 | 35.6 ± 14.9 | 36.9 ± 20.6 | 51.3 ± 17.2 | 45.2 ± 13.2 | 50.1 ± 9.9 |
| **VM (%max)** | 29.3 ± 26.2 | 35.1 ± 11.4 | 41.5 ± 15.9 | 52.3 ± 16.1 | 49.8 ± 16.5 | 52.9 ± 17.8 |
| **RF (%max)<sup>aa</sup>** | 18.5 ± 18.5**<sup>#</sup> | 40.3 ± 17.2 | 37.5 ± 11.6 | 57.2 ± 12.5 | 54.7 ± 13.3 | 53.6 ± 15.2 |

**Table 1:** Mean muscle activation of the VL, VM, and RF at the start of and during the isokinetic phase of the contraction at $100^\circ \cdot s^{-1}$ and $300^\circ \cdot s^{-1}$ .
|  | Onset of concentric contraction |  |  | Isokinetic phase |  |  |
| --- | --- | --- | --- | --- | --- | --- |
|  | NoPRE | PRE<sub>iso</sub> | PRE<sub>ecc</sub> | NoPRE | PRE<sub>iso</sub> | PRE<sub>ecc</sub> |
| **100°·s<sup>-1</sup> condition** |  |  |  |  |  |  |
| **VL (%max)<sup>a</sup>** | 9.1 ± 12.9*## | 44.6 ± 15.6## | 33.6 ± 13.5 | 43.6 ± 7.8 | 42.9 ± 10.5 | 45.7 ± 10.2 |
| **VM (%max)<sup>a</sup>** | 14.3 ± 15.8*# | 48.4 ± 19.7## | 34.1 ± 12.0 | 45.3 ± 9.8 | 43.6 ± 12.0 | 49.8 ± 14.5 |
| **RF (%max)<sup>a</sup>** | 7.4 ± 11.0*## | 41.4 ± 16.8 | 32.4 ± 11.6 | 48.9 ± 8.6 | 49.2 ± 9.6 | 50.8 ± 11.5 |
| **300°·s<sup>-1</sup> condition** |  |  |  |  |  |  |
| **VL (%max)** | 26.4 ± 26.7 | 35.6 ± 14.9 | 36.9 ± 20.6 | 51.3 ± 17.2 | 45.2 ± 13.2 | 50.1 ± 9.9 |
| **VM (%max)** | 29.3 ± 26.2 | 35.1 ± 11.4 | 41.5 ± 15.9 | 52.3 ± 16.1 | 49.8 ± 16.5 | 52.9 ± 17.8 |
| **RF (%max)<sup>aa</sup>** | 18.5 ± 18.5**# | 40.3 ± 17.2 | 37.5 ± 11.6 | 57.2 ± 12.5 | 54.7 ± 13.3 | 53.6 ± 15.2 |

#### Torque-angle and torque-length relationships modeling

At 100°.s^-1^, torque-angle and torque-length relationships in all pre-activation conditions were well fitted by the model (Figure 6, Figure 7) with very high values of R²: a mean R² of 0.94 ± 0.11 for the torque-angle relationship (from 0.92 ± 0.14 in PRE_iso_ to 0.95 ± 0.09 in NoPRE), and a mean R² of 0.93 ± 0.10 for the torque-fascicle length relationship (from 0.89 ± 0.16 in PRE_iso_ to 0.95 ± 0.06 in NoPRE). As shown in Figure 6, the torque produced during the isokinetic phase in the 300°.s^-1^ condition in PRE_iso_ and PRE_ecc_ did not exhibit an inverted U-shaped pattern. Thus, the torque-angle and torque-length relationships could not be adequately and robustly modeled for the majority of participants (see Supplementary Fig. S2), and will not be included further in the analysis.

**Figure 7:**
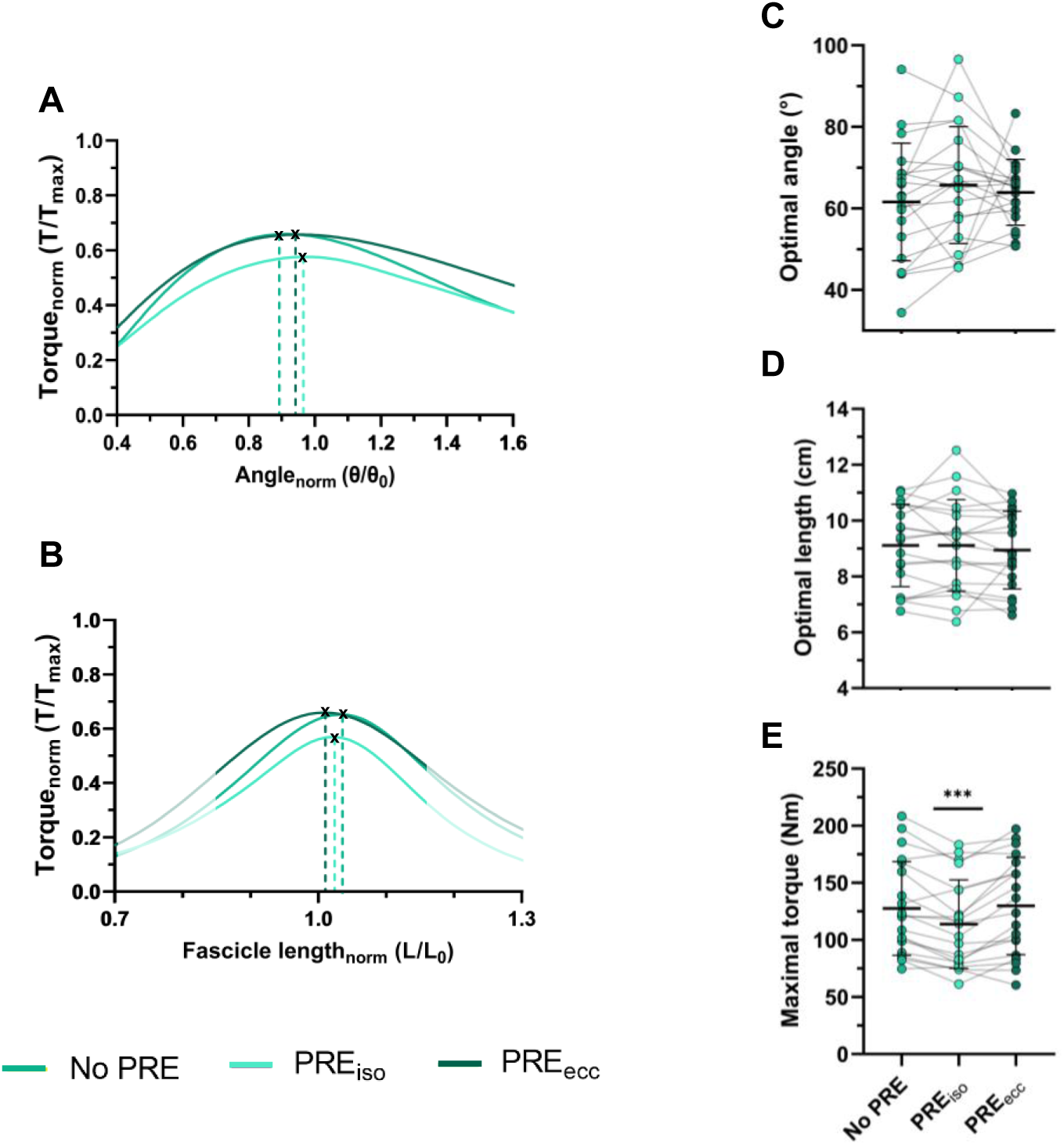
Normalized torque-angle (A) and torque-length (B) relationships in isokinetic contractions at 100°.s^-1^ preceded by no pre-activation (NoPRE), isometric pre-activation (PRE_iso_), or eccentric pre-activation (PRE_ecc_). Data in panels A and B are means; standard deviations are not shown for clarity. For panels A and B, non-shaded data correspond to the range of data observed during the experiments. Panels C, D, and E represent means and standard deviations (black) and individual data (colored dots) of the optimal angle, optimal length, and maximal torque, respectively. A main effect of pre-activation was detected for maximal torque, and *** represents a significant difference compared with NoPRE and PRE_ecc_ (p<0.001) in post-hoc analysis. N=20.

A main effect of pre-activation condition was found on T_max_ (p<0.001, Figure 7E). T_max_ was significantly lower in PRE_iso_ compared with NoPRE (p<0.001) and PRE_ecc_ (p<0.001), while no significant difference was found between NoPRE and PRE_ecc_ (p=1.000). The absolute optimal angle did not differ between the three pre-activation conditions (p=0.394, Figure 7C), with values ranging from 61.7 ± 14.6° in NoPRE to 65.7 ± 14.3° in PRE_iso_. ANOVA also revealed no effect of pre-activation on absolute optimal fascicle length (p=0.522, Figure 7D), with values ranging from 9.1 ± 1.5 cm in NoPRE to 9.0 ± 1.4 cm in PRE_ecc_.

## DISCUSSION

This study is, to our knowledge, the first to demonstrate that an increase in knee joint angular velocity is associated with some alteration of the optimal joint angle at peak torque, but not of the optimal fascicle length, partially confirming our first hypothesis. The effect of pre-activation is dependent on the contraction modality. Despite an increase in torque production during the initial phase of the concentric contraction, contrary to our second hypothesis, we did not observe an increase in mean and maximal torque during the isokinetic phase of contractions with eccentric or isometric pre-activation. We even observed a reduction in torque-generating capacity during this phase under isometric pre-activation conditions.

### Effect of joint velocity

In line with the force-velocity relationship, increasing the knee joint velocity from 50 to 300°.s^-1^ resulted in an increase in fascicle velocity, as well as a decrease in maximal torque production (Brito Fontana et al., 2014; Forrester et al., 2011; Hauraix et al., 2017). For the first time, torque-angle and torque-length relationships at high knee joint velocities (i.e. >400°.s^-1^, Figure 3) were investigated. However, it was not possible to robustly fit torque-angle and torque-length relationships above 300°.s^-1^, which may be mainly due to the flattening of torque-angle patterns with increasing joint velocity (Anderson et al., 2007). Moreover, the necessity of considering only a period of contraction under quasi-isokinetic conditions for analysis (to avoid the confounding effect of joint velocity within the relationship) resulted in a substantial reduction in the range of motion (and thus fascicle length) available for modeling both relationships. In the extreme 450°.s^-1^ condition, two factors may have reduced duration of the isokinetic phase and, consequently, the ability to reliably fit the torque-angle and torque-length relationships. First, acceleration was lower at the onset of movement compared with the other isokinetic conditions due to dynamometer settings. Second, the limited total range of motion in our study, and particularly the most extended knee angle achieved (i.e. ∼30°), may have exacerbated this limitation by triggering dynamometer deceleration earlier. Nevertheless, modeling these relationships at velocities ranging from 0° to 300°.s^-1^ provided a comprehensive overview of their alterations with increasing joint velocity.

Our results show a main effect of joint velocity on the optimal angle, with changes in the optimal angle as angular velocity increased (Figure 5). Although we failed to detect significant differences between conditions in the post-hoc analysis, the optimal angles observed at joint velocities ranging from 0°.s^-1^ to 200°.s^-1^ appear consistent with several previous studies demonstrating a shift in the optimal angle toward a more extended knee position (Bobbert & Van Ingen Schenau, 1990; Thorstensson et al., 1976). This shift in the optimal angle could be related to the time required to achieve maximal muscle activation, described as the “active state” by Bobbert and Van Igen Schenau (1990). However, in the present study, EMG activity during the isokinetic phase did not differ between conditions from 50°.s^-1^ to 300°.s^-1^, leading us to conclude that the hypothesis of reduced activation is unlikely to explain our findings. Another hypothesis suggests that the optimal fascicle length remains unchanged at high velocities compared with slow velocities, through compensation by elongation of the series elastic component (Kawakami et al., 2002). These authors reported similar changes in muscle-tendon unit behavior during isokinetic contractions and in series elastic component elongation during isometric contractions. Our results support this hypothesis, as we show that the torque-fascicle length relationship is not influenced by movement velocity. Thus, for a given joint angle, fascicle length would be greater at high velocities than at slow velocities. This is consistent with the findings of Reeves & Narici (2003), who demonstrated that increasing ankle joint velocity was associated with a small increase in operating fascicle length for a given joint angle. Our results extend these findings by showing for the first time that the optimal fascicle length is not altered by joint velocity. Consequently, changes in the optimal angle should be interpreted with caution, as they may not reflect a change in optimal fascicle length, but rather a combination of factors, such as activation dynamics or electromechanical delay. Finally, these results suggest that muscle-tendon interactions may permit a different behavior between the torque-angle and the torque-fascicle length relationships.

### Effect of pre-activation

As expected, we observed an increase in torque in the first part of the concentric contraction with pre-activation compared with NoPRE (Figures 6A and 6E), and, similarly to Goecking et al. (2024), torque did not differ between PRE_iso_ and PRE_ecc_ at either velocity. Consistent with our hypothesis and the findings of Finni et al. (2003), pre-activation resulted in greater EMG activity at the start of the concentric contraction, which could explain the higher torque observed in PRE_iso_ and PRE_ecc_ during this phase. Moreover, we observed shorter fascicle lengths in PRE_iso_ and PRE_ecc_ in the initial phase of the concentric contraction, consistent with the findings of Beaumatin et al. (2018). This shortening could reduce tendon slack (Ichinose et al., 2000) and may be more favorable for force production, as the fascicles would operate closer to the plateau of the torque-length relationship. Fascicle shortening velocity was also decreased during this phase in PRE_iso_ and PRE_ecc_, similarly to Goecking et al. (2024), thereby facilitating greater force production according to the force-velocity relationship.

However, during the isokinetic phase, we did not observe an increase in torque in PRE_ecc_ compared with NoPRE and, more surprisingly, maximal torque in PRE_iso_ was lower than that observed in NoPRE and PRE_ecc_. This finding differs from that of Narici et al. (1991), who did not report any difference in isokinetic peak torque with pre-activation. In our study, the isometric pre-activation was maximal, whereas it was performed at 25% of MVIC in the study by Narici et al. (1991). Furthermore, this discrepancy is unlikely to be explained by dynamic fascicle behavior in PRE_iso_, since fascicle length at the onset of the concentric contraction, fascicle velocity, and, more globally, the patterns of the torque-angle and torque-length relationships did not differ from those observed in PRE_ecc_ (Figures 7A and 7B). Furthermore, this finding is unlikely to be explained by tendon properties, as Holzer et al. (2023) demonstrated through simulation that MTUs with stiff tendons, such as the quadricipital tendon (Karamanidis & Arampatzis, 2006), produce similar amounts of force produced in contractions performed with and without pre-activation. Finally, because EMG activity was similar across all conditions, a deficit in neural activation is unlikely to have contributed to the torque deficit observed following isometric pre-activation.

Therefore, we speculate that the most likely explanation is the occurrence of a phenomenon known as residual force depression (rFD). Residual force depression is characterized by a reduction in maximal torque-producing capacity during a fixed-end contraction following active muscle shortening and is thought to result from the giant titin protein inhibiting cross-bridge formation within the sarcomere (for reviews, see Chen et al. 2019; Hahn et al., 2023). Although rFD is generally observed during isometric contractions preceded by active shortening (Lee & Herzog, 2003; Raiteri et al., 2024), it has also been demonstrated following isometric contractions (Goecking et al., 2024; Holt & Williams, 2018), given that fascicles shorten during isometric pre-activation (Ito et al., 1998). Because the magnitude of rFD depends on fascicle work (Lee & Herzog, 2003; Raiteri et al., 2024), it appears that the amount of fascicle shortening work performed during maximal isometric pre-activation is sufficient to induce rFD. This effect may persist throughout the subsequent isokinetic phase of the contraction, as proposed by Goecking et al. 2024 and McDaniel et al. (2010), thereby contributing to the greater reduction in torque following isometric pre-activation.

In our study, we did not observe an increase in torque in PRE_ecc_ compared with NoPRE during the isokinetic phase of the contraction, contrary to the findings of Goecking et al. (2024). This discrepancy may be explained by differences in the MTUs investigated, as we examined knee extensions, whereas Goecking et al. (2024) studied plantar flexions. As previously mentioned, tendon stiffness may influence the effect of pre-activation on torque production (Holzer et al., 2023), potentially resulting in similar torque production between PRE_ecc_ and NoPRE in our study. Likewise, we speculate that the absence of a difference between PRE_ecc_ and NoPRE may be explained by the presence of residual force enhancement (rFE), which may have attenuated the effect of rFD during the isokinetic contraction (Goecking et al., 2024). Residual force enhancement is characterized by an increase in force-producing capacity during a fixed-end contraction following active lengthening and is thought to result from an increase in passive force associated with titin (for reviews, see Fukutani & Herzog, 2019; Hessel et al., 2017). Following the eccentric pre-activation, we hypothesize that rFE may have partially offset the negative effect of rFD during the subsequent concentric contraction (Fortuna et al., 2019), resulting in no significant difference in torque between PRE_ecc_ and NoPRE.

We were unable to determine the effect of pre-activation on the optimal angle and optimal fascicle length at 300°.s^-1^ because we were unable to model the torque-angle and torque-length relationships for the majority of participants. This issue may be related to a combined effect of acceleration correction and pre-activation, where torque production at the beginning of the contraction was sufficiently high to prevent the formation of the characteristic inverted U-shaped pattern during the isokinetic phase. Nevertheless, mean torque produced during the isokinetic phase remained lower in PRE_iso_ than in PRE_ecc_ and NoPRE, confirming the findings at 100°.s^-1^ and suggesting that isometric pre-activation may be disadvantageous when the objective is to maximize torque production.

### Implications for isokinetic testing

This study highlighted that the operating fascicle length may be slightly modified at a given joint angle with increasing velocity; however, this change was not accompanied by a shift in optimal fascicle length. This finding is particularly relevant given the common assumption that the torque-angle relationship can be used as a proxy for the torque-length relationship (Kellis & Blazevich, 2022). Therefore, as previously mentioned, caution should be taken when using the torque-angle relationship to infer fascicle behavior, as alterations in the optimal angle do not necessarily reflect shifts in optimal fascicle length. Moreover, from a practical perspective, these findings suggest that the torque-length relationship can be assessed dynamically during isokinetic contractions to estimate the force-fascicle length relationship and the associated optimal fascicle length. This approach is particularly attractive for researchers because it requires less testing time and induces less fatigue than performing multiple maximal isometric contractions. In addition, this study provides important insights into the interpretation of torque production during isokinetic testing. Our results suggest that peak torque values can be used to compare force-producing capacity across different joint velocities, even when peak torque occurs at slightly different joint angles. Indeed, peak torque still appears to reflect the maximal force-generating capacity of the participants at a similar optimal fascicle length across velocities. Therefore, minor differences in optimal angle with increasing velocity do not appear to compromise the interpretation of peak torque values. Furthermore, although changes in the torque-angle relationship were observed, these alterations remained relatively small, and our statistical analyses did not allow us to determine precisely at which velocity they emerged. Alternatively, torque can be analyzed using mean values over a range of motion. In this case, our findings emphasize the importance of selecting a sufficiently large range of motion (approximately 30-50°) centered within the functional range of motion to ensure relatively stable angular velocity conditions across all tested velocities. Under these circumstances, using the same angular region across velocities is unlikely to induce substantial bias into the analysis. Nevertheless, our results also highlight the difficulty of modeling torque-angle and torque-length relationships above 300°.s^-1^. This limitation is partly attributable to the flattening of the torque profile at high velocities, but also to dynamometer constraints that limited the ability to maintain the target velocity throughout a sufficient range of motion. One potential solution would be to model the three-dimensional torque-length-velocity relationship, thereby accounting for both inertial effects and dynamometer related constraints.

Several studies have recommend performing isokinetic contractions with pre-activation (Brown et al., 2016; Forrester et al., 2011; Germano Maciel et al., 2020; Pain et al., 2013). The primary rationale for including pre-activation is to ensure full muscle activation from the onset of the isokinetic contraction and to reduce tendon slack, thereby maximizing torque production. Although our results confirmed greater EMG activity and higher torque production at the onset of the concentric contraction with both isometric and eccentric pre-activation (Figure 5), EMG activity during the subsequent isokinetic phase was similar across conditions. Moreover, maximal torque did not differ between NoPRE and PRE_ecc_, whereas PRE_iso_ resulted in lower torque production than the other conditions. Therefore, pre-activation appears beneficial only when maximizing torque production at the beginning of the range of motion is required, regardless of contraction velocity. In this context, maximal eccentric pre-activation appears to be the most appropriate strategy because it allows participants to produce greater torque from the onset of the concentric contraction to the end. Consequently, eccentric pre-activation may increase total positive work performed over the entire range of motion compared with NoPRE or PRE_iso_, which could be advantageous in training or intervention protocol. Importantly, eccentric pre-activation did not alter peak torque or mean torque during the isokinetic phase compared with the NoPRE condition. In contrast, isometric pre-activation resulted in an underestimation of torque production, affecting both mean torque during the isokinetic phase and peak torque values. Accordingly, we recommend that future studies using isokinetic contractions avoid maximal isometric pre-activation and instead use either a conventional contraction protocol or eccentric pre-activation.

## CONCLUSION

The present study demonstrated that, despite alterations in the optimal joint angle with increasing joint velocity, the torque-fascicle length relationship remained unchanged, with a similar optimal fascicle length across conditions. This finding, suggests that shifts in the torque-angle relationship should not be interpreted as alterations in the intrinsic force-length properties of the muscle. Future experimental studies should further investigate this phenomenon by examining muscle-tendon units with more compliant tendons, such as the gastrocnemius medialis, to determine whether larger shifts in the torque-angle relationship occur without corresponding changes in the torque-fascicle length relationship. In addition, our results indicate that using maximal isometric pre-activation before an isokinetic contraction is not recommended, as it reduced maximal torque production compared with eccentric pre-activation or no pre-activation, at both slow and fast velocities. These findings have important implications for future investigations of torque-length relationships and, more broadly, for researchers and practitioners conducting isokinetic assessments. Finally, future studies should seek to better understand the influence of contraction history during dynamic contractions, particularly the potential roles of transient rFE and rFD. A clearer understanding of these phenomena would provide valuable insights into the mechanisms governing torque production during isokinetic contractions.

## SUPPLEMENTARY MATERIAL

Supplementary Fig. S1: https://doi.org/10.57745/BLSDIH

Supplementary Fig. S2: https://doi.org/10.57745/BLSDIH

## DATA AVAILABILITY

Datasets generated during the current study are available following this link: https://doi.org/10.57745/BLSDIH

## ACKNOWLEDGMENTS

Preprint is available on Biorxiv:

## AUTHOR CONTRIBUTIONS

TT, AN and SD conceived and designed research, TT and LL performed experiments, TT analysed data, TT, AN and SD interpreted results of experiments, TT prepared figures, TT drafted manuscript, TT, AN and SD edited and revised manuscript, TT, AN, LL and SD approved final version of manuscript.

